# Influence of Substrate Elastic Modulus on Cell Shape and Displacement During the Motility Cycle

**DOI:** 10.1101/2025.05.26.656039

**Authors:** Arata Nagai, Hiromu Kuwabara, Yuuta Moriyama, Toshiyuki Mitsui

## Abstract

Cardiac fibroblasts play a pivotal role in heart tissue dynamics, responding to mechanical cues from their environment. This study examines how substrate elasticity influences the motility and shape of chicken embryonic cardiac fibroblasts on polydimethylsiloxane (PDMS) substrates of different stiffness (10 kPa and 400 kPa). Using time-lapse phase-contrast microscopy, we identified distinct motility cycles characterized by elongation, contraction, and retraction phases. On soft substrates, cells exhibited slower retraction speeds and longer motility cycles compared to those on hard substrates. Notably, cells on soft substrates left trailing edges upon retraction, leading to pronounced morphological oscillations, whereas cells on hard substrates adopted near-circular shapes with less coordinated motility. Quantitative analyses revealed that the duration of the static phase was significantly shorter on hard substrates, correlating with differences in cell shape parameters, including length, circularity, and area. These findings suggest that substrate elasticity modulates the balance between cell-substrate adhesion and contractile forces, influencing cell polarity and motility. Our work advances the understanding of how mechanical environments shape fibroblast dynamics, offering insights relevant to tissue engineering and fibrosis research.

**SIGNIFICANCE:** Mechanical cues from the cellular microenvironment are known to regulate cell behavior, yet how substrate stiffness shapes the dynamic motility cycle of fibroblasts remains incompletely understood. Here, we use high-resolution time-lapse imaging to reveal that cardiac fibroblasts on soft (10 kPa) PDMS undergo polarized, oscillatory migration with prolonged static phases, while those on stiff (400 kPa) substrates exhibit rapid retraction and random-like displacement with reduced polarity. These findings highlight how substrate elasticity modulates the interplay between traction forces and adhesion stability, governing directionality and morphology. This work provides mechanobiological insights into fibroblast behavior with implications for designing biomaterials to guide tissue remodeling and fibrosis progression.

## INTORDUCTION

The intricate dynamics of cellular behavior in response to the physical environment have become a focal point in tissue engineering and regenerative medicine (1). In the heart, fibroblasts—often overshadowed by cardiomyocytes— play a pivotal role in tissue maintenance and repair by responding to mechanical cues (2). Under physiological conditions, cardiac fibroblasts regulate extracellular matrix (ECM) remodeling to support normal development and wound healing (3,4). However, in pathological states, prolonged mechanical stress can induce fibroblast-to-myofibroblast transition, leading to fibrosis and adverse cardiac remodeling, ultimately contributing to heart disease (2,5).

Recent advances in bioengineering have emphasized the importance of designing soft materials that closely replicate in vivo environments to enable precise in vitro investigations of cellular behavior (6,7). Among these, polydimethylsiloxane (PDMS) and hydrogels with tunable mechanical properties have emerged as critical tools for studying mechanobiological responses. Modulating substrate stiffness, topography, and tension allows researchers to investigate how cells adapt in terms of migration, proliferation, and differentiation (8,9). In particular, single-cell studies suggest that cellular motility is largely dictated by ECM stiffness and external mechanical factors (10,11). Additionally, cell migration speed is regulated by adhesion strength to the substrate, highlighting the importance of mechanical interactions in governing cell motility (12).

External mechanical factors influencing cell motility are highly dependent on cell type and ECM stiffness (13). For example, neuronal cells migrate more efficiently on soft matrices (∼50 kPa) resembling brain tissue, while fibroblasts and muscle cells exhibit motility on stiffer substrates (10–100 kPa), mimicking muscle and fibrotic tissues, including the heart (14-16). In fibroblasts, motility is also significantly influenced by adhesion strength, which is modulated by extracellular protein coatings on the substrate (17). Even when substrate stiffness remains constant, variations in fibronectin or collagen coating concentrations alter adhesion stability, ultimately affecting migration behavior (18). These findings highlight the intricate interplay between substrate properties, adhesion dynamics, and cell morphology in regulating motility patterns.

PDMS, with its tunable stiffness and surface properties, has become a versatile material for in vitro studies, providing a controlled environment to investigate fibroblast behavior (19). The flexibility of PDMS in mimicking the mechanical range of soft tissues makes it particularly effective in preserving fibroblast phenotypes and studying their mechanobiological responses (20-23). On the other hand, increased substrate stiffness and enhanced adhesion due to extracellular protein coatings have been shown to promote fibroblast differentiation into myofibroblasts (20,23). To further explore this mechanobiological response, we used PDMS substrates with stiffnesses differing by an order of magnitude—10 kPa and 400 kPa—representing two extremes commonly tested in fibroblast mechanobiology studies (10,24,25). By applying coatings that reduced adhesion strength, we created weaker attachment conditions and observed the dynamics of isolated single cells over extended durations with high temporal resolution. Our findings reveal that detachment-induced changes in cell shape significantly influence both the directionality and speed of fibroblast migration.

To analyze cellular motility, we examined temporal variations in cell displacement and centroid position, along with structural transformations occurring over a 24-hour period. Typically, single-cell migration on a substrate follows a cyclical process consisting of three phases: an initial extension phase, where one end elongates to establish new adhesions; a static elongated phase; and a retraction phase, which shifts the cell centroid. While the phenotypic and molecular mechanisms underlying these transitions have been extensively studied, the precise biophysical movements and morphological alterations within each phase have received less attention. Previous studies have generally described migration in terms of broad movement cycles rather than the detailed transitions within each cycle. In this study, we specifically analyzed the dynamics of each phase and discovered that post-detachment cell morphology significantly dictates the directionality of subsequent movement. Our findings offer novel insights into the interplay between mechanical and adhesive cues in regulating fibroblast motility. Understanding these dynamics is crucial for uncovering the mechanobiological pathways governing fibroblast behavior and their potential implications for tissue remodeling and fibrosis.

## RESULTS

We studied the influence of substrate elasticity on cellular morphology and motility by seeding cardiac fibroblasts on 10 kPa and 400 kPa PDMS substrates under weaker adhesion conditions. The dynamics of individual cells were captured using time-lapse microscopy at Δt = 5 minutes over a 24-hour period, focusing on isolated cells to ensure intrinsic behavior (26). Observations exceeding 24 hours revealed distinct cycles of characteristic cellular motions (12,27). Each motility cycle begins with cell extension, characterized by the formation of a protrusion at one end and the development of new adhesions at the leading edge while elongating its shape (28). This is followed by a static phase, during which the cell maintains its shape and generates polarity increasing contractile force (29). The cycle then proceeds to retraction, where the rear of the cell detaches and relaxes, facilitating forward motion and completing the cycle before the next motility cycle begins.

### Motility Patterns on 10 kPa Substrate

On 10 kPa PDMS, fibroblasts exhibited oscillatory behavior with coordinated back- and-forth displacements as part of a repeated motility cycle (Figs. 1A–1D). Each figure (a–d) illustrates characteristic cell shape changes corresponding to the steps in the motility cycle. For example, in Group A (t = 0–80 minutes) shown in Fig. 1A, the cell initiates protrusion formation (arrowhead, Fig. 1Ab) in the direction indicated by the red arrow, followed by elongation to 71 µm. During the static phase, the trailing edge thins (arrowhead, Fig. 1Ac), and within 10 minutes, the cell retracts, reducing its length to 51 µm while moving forward (Fig. 1Ad). The centroid trajectory (Fig. 1Ae) confirms directional displacement along the Y-axis, capturing one complete motility cycle.

**FIGURE 1.**
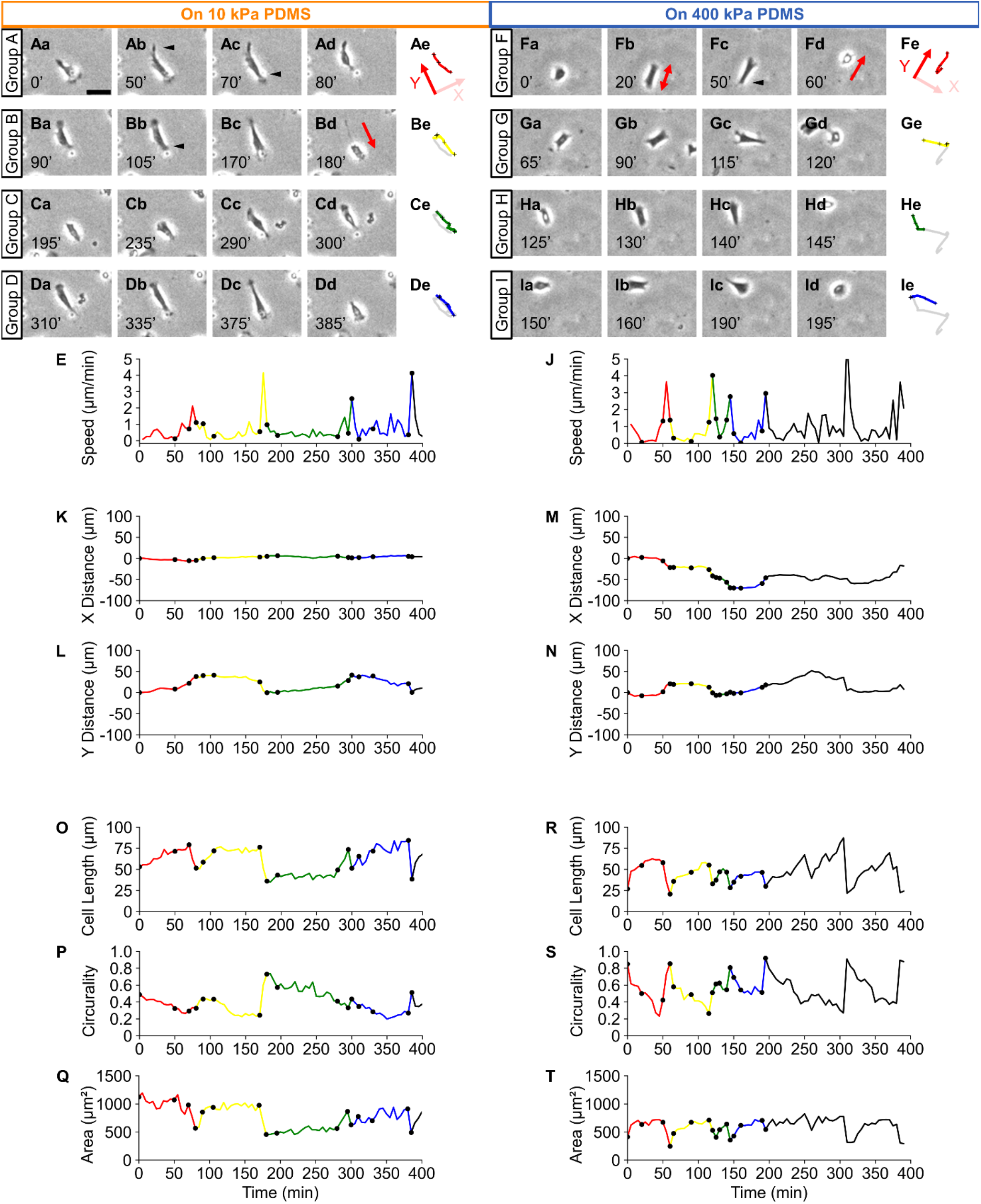
Periodic shape changes and motility cycles of cardiac fibroblasts on 10 kPa and 400 kPa PDMS substrates. (Aa–d, Ba–d, Ca–d, Da–d) Phase-contrast images of a single cell on 10 kPa PDMS. (Ae, Be, Ce, and De) Cell trajectories for Groups A–D. Arrows indicate elongation directions (red), retraction directions (yellow), and arrowheads mark protrusions and trailing edges. Scale bar: 50 µm. (E) Cell speed on 10 kPa PDMS. (Fa–d, Ga–d, Ha–d, Ia–d) Phase-contrast images of a single cell on 400 kPa PDMS. (Fe, Ge, He, and Ie) Cell trajectories for Groups F–I. (J) Cell speed on 400 kPa PDMS. (K–L) X- and Y-axis centroid displacements on 10 kPa PDMS. (M–N) X- and Y-axis centroid displacements on 400 kPa PDMS. (O–Q) Cell morphology on 10 kPa PDMS: cell length (O), circularity (P), and area (Q). (R–T) Cell morphology on 400 kPa PDMS: cell length (R), circularity (S), and area (T). Plot line colors correspond to Groups A–D (10 kPa) and Groups F–I (400 kPa). Black dots indicate the times when phase-contrast images were taken: Aa– Dd (10 kPa) and Fa–Id (400 kPa).

In Group B (t = 80–180 minutes), as shown in Figs. 1Ba–1Bb, the cell reverses its direction, forming a new protrusion at the previously trailing edge (arrowhead). The cell elongates to 76 µm (Fig. 1Bc) before retracting within 10 minutes, shortening to 36 µm, and migrating back toward its original position (red arrow, Fig. 1Bd). The trajectory in Fig. 1Be confirms this back-and-forth motion. Similar behavior was observed in Groups C and D, with each cycle consisting of an extension phase, a static phase lasting approximately 1 hour, and ending with rapid retraction (Fig. 1Ce, 1De). As a result, each complete motility cycle lasted nearly 100 minutes with the centroid showing oscillatory displacement along a single line (Fig. 1De). These dynamic behaviors are detailed in Supplementary Movie SI 1. Speed analysis from centroid displacement (Fig. 1E) revealed peaks exceeding 2 µm/min during retraction phases, while speeds remained below 1 µm/min (12,30) during gradual protrusion and adhesion phases. Given that the typical speed for cellular motion is below 1 µm/min (30), the rapid shrinkage observed during trailing-edge detachment constitutes a notable finding.

### Motility Patterns on 400 kPa Substrate

On 400 kPa PDMS, fibroblasts exhibited anisotropic back-and-forth displacements relative to the 10 kPa substrate, resembling random-walk-like behavior. As shown in images a to d of Groups F through I in Figs. 1F, 1G, 1H, and 1I, the cells underwent repeated protrusion-retraction cycles. The cell trajectories (Figs. 1Fe to 1Ie) lacked directional regularity throughout the cycle.

For example, in Group F (t = 0–50 minutes), the cell extended protrusions toward both edge ends (red arrows, Fig. 1Fb). During the static phase, the cell maintained nearly length of 55 µm (Figs. 1Fb–1Fc), while one edge began to thin (arrowhead, Fig. 1Fc), showing cell polarity. The duration of the static phase was shorter than that observed on the 10 kPa substrate. Within 10 minutes, the cell retracted to 21 µm, forming a nearly circular shape (Fig. 1Fd). Similar cell behaviors were observed in Groups G, H, and I, with cells exhibiting static phases followed by rapid retraction. The static phase durations were notably shorter on the 400 kPa substrate compared to the 10 kPa substrate. Directional shifts and random-like oscillatory activities were observed in these groups, as shown in Figs. 1F–1I, with Fig. 1Ie illustrating this random-like displacement pattern.

Speed analysis (Fig. 1J) revealed peaks exceeding 2 µm/min during retraction phases, while speeds during protrusion remained below 1 µm/min (12,30). These speed peaks signified the onset of retraction phases within the motility cycle. The overall cycle period on the 400 kPa substrate was estimated to be approximately 50 minutes, shorter than the nearly 100-minute cycles observed on the 10 kPa substrate. These dynamic behaviors are visually detailed in Supplementary Movie S2.

### Effect of Substrate Elasticity on Directionality of Cell Displacement

To investigate how substrate elasticity influences the directionality of cell motion, cell displacement was decomposed into X and Y components, as shown in Fig. 1Ae for 10 kPa and Fig. 1Fe for 400 kPa, and analyzed in Figs. 1K, 1L and Figs. 1M, 1N, respectively. For 10 kPa, Fig. 1L illustrates oscillatory displacements along the Y-axis with an amplitude of approximately 50 µm during retraction phases, with cell speeds exceeding 2 µm/min. Displacement along the X-axis remained confined within ± 5 µm. Conversely, at 400 kPa, cells exhibited displacements exceeding 20 µm in both X and Y directions, as shown in Figs. 1M and 1N, reflecting a more diffusive and less aligned motion pattern.

### Cell Shape During Motility Cycle on 10 kPa and 400 kPa PDMS Substrates

We analyzed cell morphology, including cell length, circularity, and area, as shown in Figs. 1O–1Q for 10 kPa and Figs. 1R–1T for 400 kPa. On 10 kPa PDMS, cell length oscillated between approximately 30 µm and 75 µm with a periodicity of about 90 minutes (Fig. 1O). Sharp reductions in length were synchronized with speed peaks during retraction phases. Throughout these oscillations, circularity remained near 0.5, as shown in (Fig. 1P), indicating an elongated shape, while the area stabilized at approximately 1000 µm^2^ during static phases (Fig. 1Q). On 400 kPa PDMS, cells predominantly maintained lengths below 50 µm, with retraction phases causing sharp reductions to less than 30 µm (Fig. 1R). As a result, the cell shape became nearly circular, with circularity approaching 1.0 (Fig. 1S). Concurrently, cell area decreased to a minimum during retraction phases before recovering to slightly over 500 µm^2^ during static phases (Fig. 1T).

### Statistical Analyses of Directionality of Cell Velocity and Cell Motility Cycle on 10 kPa and 400 kPa PDMS Substrates

From the correlation between cell motility on both PDMS substrates, it was evident that differences in directionality and periods of the motility cycle exist. To investigate these differences statistically, we analyzed directionality and motility cycle periods on 10 kPa and 400 kPa PDMS substrates. For directionality, the angular distribution histogram of cell velocity during the elongation and retraction phases (Fig. S1A) showed linear motion for isolated cells (n = 4) on 10 kPa PDMS, as illustrated in Figs. S1A–S1D. In contrast, the angular distribution histogram of cell velocity on 400 kPa PDMS (Figs. S1E–S1H) indicated isotropic displacements during motility cycles (n = 4), suggesting a lack of directional alignment on the stiffer substrate.

Regarding the motility cycle period, cells on the 10 kPa substrate exhibited significantly longer cycles, with an average duration of 79.64 ± 7.53 minutes (n = 14), compared to 37.01 ± 2.47 minutes (n = 72) on the 400 kPa substrate (Fig. S2).

### Cell Shape transition and the Statistical Analyses of duration of Phases in motility Cell cycle, cell length, circularity and Area

To further investigate the influence of substrate elasticity on the timescales of extension, static, and retraction phases, we analyzed shape deformation during one motility cycle at 5-minute temporal resolution. Representative phase-contrast time-lapse images are shown in Figs. 2Aa–2Ae for 10 kPa and Figs. 2Ba–2Be for 400 kPa. On the 10 kPa substrate, the cell displayed pronounced anisotropic elongation with a distinct fan-shaped lamellipodium oriented in the direction of motion, while a trailing edge was retained on the opposite side (Fig. 2Aa). Retraction detached the trailing edge (arrowhead, Fig. 2Ab), generating displacement. The cell then reversed polarity by forming a new fan-shaped lamellipodium at the previous trailing edge, enabling oscillatory motion (Figs. 2Ac–2Ad). On the 400 kPa substrate, the cell exhibited a shorter fan-shaped lamellipodium. Following retraction (arrow, Fig. 2Ba), the cell adopted a circular shape with reduced length, showing less defined directionality (Fig. 2Bb). While the cell elongated (Fig. 2Bc), its alignment deviated from previous displacements and initially lacked a distinct fan-shaped lamellipodium. A slight lamellipodium formed later, followed by minor displacement (Fig. 2Bd). Retraction occurred before full elongation, resulting in a circular shape and reduced polarization.

**FIGURE 2.**
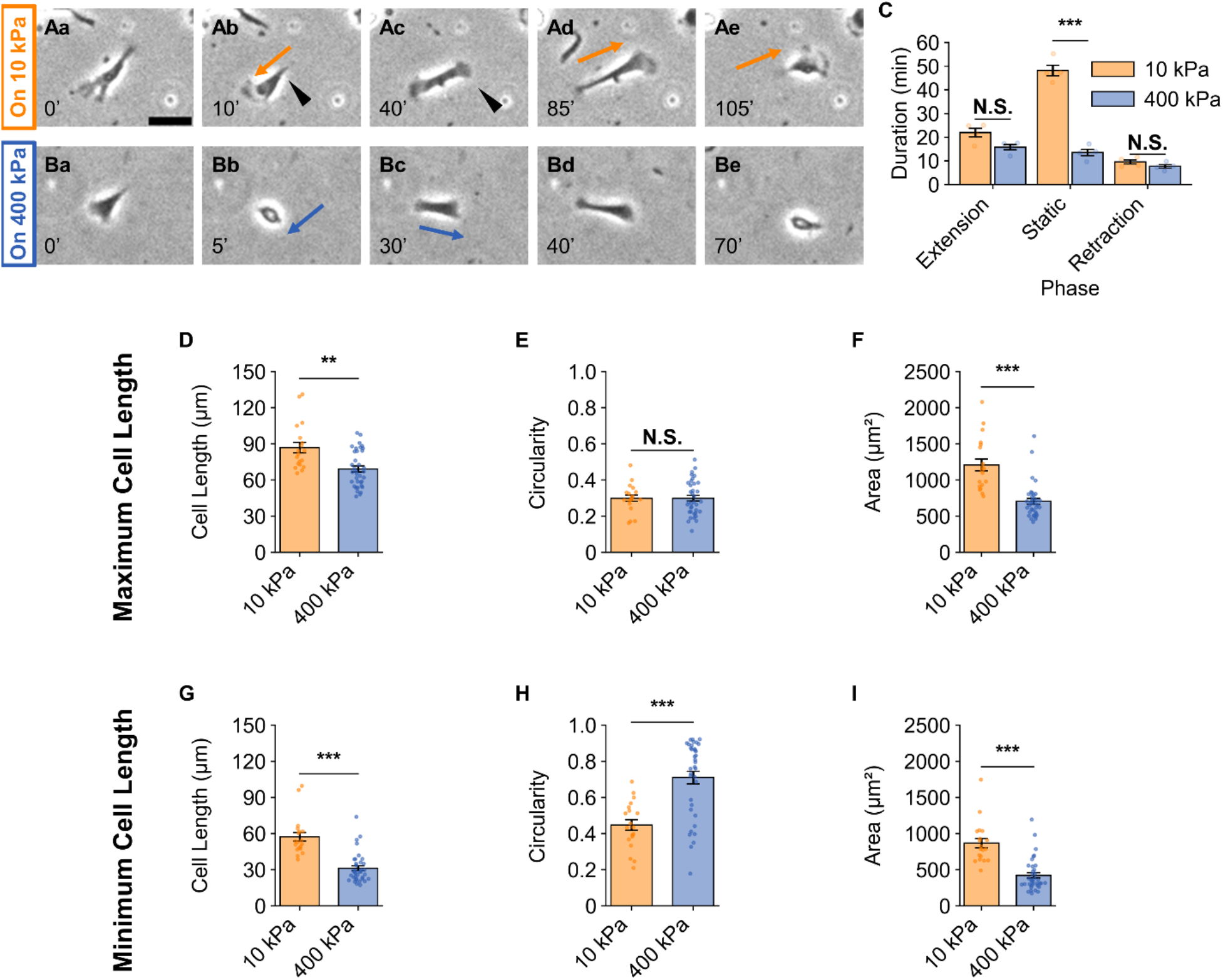
Cell shape transitions across different s on 10 kPa and 400 kPa PDMS substrates. (A, B) -contrast images of fibroblasts during motility cycles on 10 kPa (A) and 400 kPa (B) PDMS substrates. Scale bar: 50 µm. (C) Box plot of durations: Extension (1), static (2), and retraction (3) on 10 kPa and 400 kPa PDMS (n = 4 for both substrates). The static duration was significantly longer on 10 kPa PDMS compared to 400 kPa. (D– F) Box plots of cell morphology during the static on 10 kPa and 400 kPa PDMS substrates: (D) Cell length (p = 1.5797e-03), (E) Cell Circularity (p = 9.9860e-01), (F) Area (p = 1.1665e-05). (G–I) Box plots of cell morphology at the minimum cell size after the retraction on 10 kPa and 400 kPa PDMS substrates: (G) Cell length (p = 1.0332e-06), (H) Circularity (p = 6.5137e-07), (I) Area (p = 2.3187e-06). n = 19 for 10 kPa and n = 36 for 400 kPa. Data are presented as mean ± standard error. ***P < 0.001; **P < 0.01; *P < 0.05 using Welch’s t-test. N.S., not significant. n=19 for 10 kPa and n=36 for 400 kPa.

The statistical analysis of the durations of the extension, static, and retraction phases was performed using phase classification based on the ratio of elongation and cell length. The extension phase was defined as the period from the minimum cell length to elongation, where the ratio of elongation exceeded 0.1, calculated at 5-minute intervals. The static phase was identified when the elongation ratio remained below 0.1 for 5 minutes. Retraction began when the elongation ratio became negative and continued until the cell length reached its minimum.

Fig. 2C reveals that the static phase duration in isolated single cells was significantly longer on the 10 kPa substrate, averaging 57.82 ± 1.16 minutes (n = 4), compared to 21 ± 1.30 minutes (n = 4) on the 400 kPa substrate. The durations of the extension and retraction phases showed no significant differences between the two substrates. These findings suggest that the overall difference in motility cycle periods between the 10 kPa and 400 kPa substrates is primarily due to the prolonged static phase observed on the 10 kPa substrate (Fig. S2).

Additionally, we analyzed cell shape at the end of the extension phase, when cell length was nearly at its maximum, and at the minimum cell length at the end of the retraction phase, where notable differences were observed (Fig. 1). Cell length (Fig. 2D), circularity (Fig. 2E), and area (Fig. 2F) were plotted for the end of the extension phase. Similarly, cell length (Fig. 2G), circularity (Fig. 2H), and area (Fig. 2I) were analyzed for the end of the retraction phase. Cells on the 10 kPa substrate were longer compared to those on the 400 kPa substrate at both phases, corresponding to larger cell areas (Fig. 2F and 2I). However, circularity was significantly higher, approaching 1, for cells on the 400 kPa substrate, reflecting a lack of prominent polarity in cell shape. Notably, the difference in cell length between the maximum and minimum values was comparable for both substrates. Since the duration of the extension phase was similar, the extension speed was nearly identical between cells on the 10 kPa and 400 kPa substrates.

## DISCUSSION

Our study examined the influence of substrate elasticity on cellular morphology and motility by analyzing cardiac fibroblasts on 10 kPa and 400 kPa PDMS substrates. Cells on the softer 10 kPa substrate exhibited coordinated oscillatory motion with pronounced elongation and distinct fan-shaped lamellipodia, while cells on the stiffer 400 kPa substrate displayed random-walk-like motion, reduced polarization, and shorter lamellipodia. Statistical analysis revealed that the prolonged static phase on 10 kPa was the primary contributor to its longer motility cycle compared to 400 kPa. Morphological differences also reflect the role of substrate elasticity. On the 10 kPa substrate, cells extended further and retained polarity after retraction, whereas on the 400 kPa substrate, cells became more circular after retraction, exhibiting a non-directional morphology. These results emphasize the essential role of substrate elasticity in shaping cell motility behavior and morphology.

### Swift displacement after detachment

Previous studies have suggested that cellular morphology and motility are primarily influenced by substrate adhesion and elasticity, which regulate the number and area density of focal adhesions (FAs) (8,9). Additionally, FA lifetimes generally last tens of minutes, and the balance between FA formation and annihilation, along with internal cell tension, determines the dynamic state of the cell (23). Notably, high-density collagen substrates enhance FA attachment to the substrate, increasing the residence time of FAK within FAs by approximately fourfold compared to low-density collagen (31). Strong adhesion to the substrate has been suggested to enhance cell spreading (9), which in turn may suppress the cell speed. In contrast, on our PDMS substrates with reduced substrate attachment, we observed significant shape changes and rapid detachment motion in fibroblasts. This suggests that weakened adhesion contributes to swift displacement dynamics upon detachment.

### Cell shape change in motility cycle on 10 kPa and 400 kPa

Fibroblasts on stiffer substrates generally elongate more after polarization, exerting greater traction forces (32-35). In our study, cells on the 10 kPa substrate extended to an area of approximately 1200 µm^2^, while those on the 400 kPa substrate extended to about 700 µm^2^, with both conditions exhibiting similar circularity values (Fig. 2F). The smaller cell area observed on the 400 kPa substrate, approximately half of the values reported in earlier studies (36), suggests that cells on the stiffer substrate were unable to elongate as extensively. Instead, they transitioned prematurely into the retraction phase, reducing overall elongation. Traction force, an internal pulling force correlated with FA formation, plays a crucial role in cell shape dynamics (37). During the process of increasing contractile force following polarization, weaker adhesion on the 400 kPa substrate may have been insufficient to sustain FA attachment, leading to premature detachment before maximum elongation. In contrast, on softer substrates, FA maturation may proceed more effectively, allowing cells to sustain elongation for a longer duration before retraction. Since internal traction forces are higher on stiffer substrates, the imbalance between traction and adhesion results in a more circular shape on the 400 kPa substrate compared to the 10 kPa substrate (38). This mechanical instability likely contributes to the shorter duration of the static phase on the stiffer substrate, as cells detach and transition into a new motility cycle more quickly. These findings emphasize the role of substrate elasticity in stabilizing elongation and regulating the duration of motility phases, highlighting the mechanical feedback governing fibroblast migration.

### Oscillatory motion vs. random-walk-like motion induced by cell shape after retraction

The directionality of migrating cells is governed by a combination of biochemical anisotropy and intracellular signaling, involving Rho-family GTPases such as Rac, Rho, and Cdc42 (41). These molecules regulate protrusion and contraction in distinct regions of the cell, establishing front-rear polarity, which in turn determines the direction of movement. Rac is localized at the leading edge, where it promotes the formation of lamellipodia and actin polymerization through Arp2/3 complex activation. Rho, on the other hand, is enhanced at the rear, stimulating actomyosin contractility and facilitating retraction. Cdc42 plays a crucial role in coordinating Rac activity and establishing front-back polarity. Fibroblasts establish such polarization more slowly than other cell types, typically taking 30–50 mins, and maintain directional persistence as long as the intracellular concentration gradients remain unchanged (42).

In our observations, cells migrating directionally on 10 kPa substrates exhibited a well-defined lamellipodium-trailing edge. However, on the 400 kPa substrate, cells elongated bidirectionally before fully establishing polarity, indicating a failure to generate stable directional movement. This phenomenon may be attributed to weaker adhesion in combination with elevated internal traction forces, leading to premature retraction before polarity is firmly established due to positive feedback, wherein molecular signaling pathways reinforce localized accumulation of key regulatory molecules (43,44). Furthermore, cell shape became increasingly circular following retraction, as shown in Fig. 2h. Under such conditions, the cell must reestablish polarity from a non-directional state, which may contribute to the random-walk-like behavior observed in Fig. S1E-1H.

In contrast, cells on 10 kPa substrates exhibited back-and-forth motion, indicative of oscillatory movement, as shown in Figs. 2Ab, 2Ae, and Figs. S2A–S2D. This suggests that cells initially establish polarity but subsequently undergo polarity reversal following the static phase. Such reversals likely arise from redistribution of Rac, Rho, and Cdc42, wherein Rac activity declines at the front or Rho activity becomes dominant at the rear (45), leading to retraction at the leading edge and a subsequent reversal in migration direction. A potential explanation for this behavior is the dynamic balance between internal traction forces and adhesion-mediated anchoring via focal adhesions (FAs). In the absence of external directional cues, such as chemical or mechanical gradients that induce directed migration (e.g., chemotaxis or durotaxis), intracellular molecular concentrations may undergo spontaneous oscillations, leading to periodic front-to-rear polarity switching. Oscillatory migration has observed in mesenchymal cells and the mechanism is proposed to result from sustained biochemical cycling of signaling molecules, yet the precise regulatory pathways remain under investigation (46). Nevertheless, such oscillatory cell motion has been captured in numerical simulations using reaction-diffusion models with closed boundary conditions (47).

Importantly, these discoveries were enabled by observing cells across multiple cycles with high temporal and spatial resolution under weaker adhesion conditions on substrates with order-of-magnitude differences in elasticity. Additionally, our study highlights the need to target specific cell morphologies when investigating the mechanical responses of cells to external environments (48).

## CONCLUSIONS

Our study demonstrates that substrate elasticity plays a crucial role in regulating the motility and morphology of cardiac fibroblasts. By analyzing fibroblast behavior on PDMS substrates of different stiffness (10 kPa vs. 400 kPa), we identified distinct motility patterns. Cells on the softer 10 kPa substrate exhibited oscillatory motion, characterized by static elongation phases, directional persistence, and periodic polarity reversals. In contrast, cells on the stiffer 400 kPa substrate displayed random-walk-like behavior, reduced polarity, and frequent transitions into a circular shape, likely due to shorter elongation times and weaker adhesion. These findings suggest that fibroblast motility is governed by the interplay between traction forces and substrate adhesion. On the softer substrate, stable focal adhesion maturation allowed cells to sustain elongation, leading to longer motility cycles. In contrast, weaker adhesion and higher traction forces on the stiffer substrate promoted premature retraction, resulting in rapid polarity loss and non-directional displacement. The transition into a circular shape following retraction further disrupted polarity, contributing to isotropic movement.

Our results suggest that the importance of high-temporal-resolution observations to capture phase-specific transitions within motility cycles. Future studies should explore the molecular pathways regulating cell shape transitions and adhesion dynamics. By deepening our understanding of fibroblast mechanobiology, these insights may inform tissue engineering, regenerative medicine, and the development of biomaterials designed to control fibroblast behavior in therapeutic applications.

## METHODS

### PREPARATION OF SUBSTRATE

To observe cell dynamics in environments with different Young’s moduls, polydimethylsiloxane (PDMS) was chosen as the substrate, and Sylgard-184 and CY52-276A/B were selected as the liquid silicones. To make a hard substrate, a 200 mg aliquot of pre-polymer base and a 20 mg aliquot of crosslinking curing agent were mixed. A 15 mg of this mixture was evenly spread on a 10 mm diameter No.0 coverslip (P35G-0-10-C, MATTEK) in a 35 mm dish. The silicone mixture on the coverslip was cured at 80°C for 30 minutes, and the Young’s modulus was estimated to be approximately 400 kPa (49). To make a soft substrate, a 100 mg aliquot of CY52-276A and a 100 mg aliquot of CY52-276B were mixed. A 15 mg of this mixture was evenly spread on a 10 mm diameter No.0 coverslip (P35G-0-10-C, MATTEK) in a 35 mm dish. The silicone mixture on the coverslip was cured at room temperature for 2 days, and the Young’s modulus was estimated to be approximately 10 kPa (50). For cell adhesion on PDMS, substrates were coated with 300 µL of 0.3 mg/mL collagen I-C (Nitta Zeratin, 631-00771) and incubated at room temperature for 4 hours. The coating solution was then removed, and the substrates were gently rinsed with 1× PBS (5913, Nissui) without allowing the surface to dry.

### Cell Culture

Primary cultures of cardiac cells were established, and a co-culture system was created (mainly cardiomyocytes and fibroblasts). The hearts were extracted from chicken embryos at HH stage 28, and blood vessels were removed. They were then fragmented into approximately 1 mm^3^ tissue pieces using pruning shears, and were immersed for 10 minutes in 1 mL of 3.5% trypsin (15090046, Thermo Fisher Scientific) at room temperature. Isolated cells were pressurized by 7 pipetting 31 G needles (NN-3138R, TERUMO) and subjected to two rounds of centrifugation (15,000 rpm for 5min) and 1 mL 7 pipetting, then passed through a 100 µm cell strainer (353360, CORNING). The cell suspension was aliquoted, and cells were seeded on substrata at a low cell density. Cells were kept at 37°C in humidified air containing 5% CO3 and grown in Dulbecco’s Modified Eagle Medium (DMEM)/F-13 medium (10565018, Thermo Fisher Scientific) with 100 U/mL penicillin, 100 µg/mL streptomycin (168-33191, FUJIFILM Wako) and 10% fetal bovine serum (FBS) (S1400-500, Biowest). To allow for isotropic cell adhesion, the cells were cultured on a rocking shaker for 18 hours and then left undisturbed for 3 hours before the observation was started.

### Time-Lapse Observation and Analysis of cell morphological transition in cell motility cycle

Images were captured using a phase-contrast microscope BZ-X810 (KEYENCE) equipped with a 4× objective lens (UPlanFl, N.A. 0.13, OLYMPUS), in TIFF format at 1920 × 1440 pixels (approximately 10 mm^2^) at 5 minutes intervals for over 24 hours. On the PDMS coated with collagen I-C, there were approximately 500 cells per image, and isolated fibroblasts that were not in contact with other cells were selected for analysis.

The shape and centroid of single cells were determined by analyzing contrast differences between the cytoplasm and the background using ImageJ. To examine the angular distribution of cell velocity, only velocities exceeding 0.1 µm/min were included, excluding cells in the static phase where apparent displacement was not detectable.

To define the phase durations in a cell motility cycle, the ratio of elongation was calculated as the rate of change in cell length using the following equation:

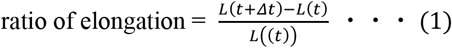

The phase classification was determined based on the ratio of elongation in the cell motility cycle: Extension (>0.1), Static (−0.06 to 0.1), and Retraction (<−0.06).

## SUPPLEMENTARY MATERIAL

See Supplementary Material for additional figures and movies illustrating the dynamics of cardiac fibroblasts on PDMS. This includes time-lapse videos of fibroblasts during some motility cycles.

## ACKNOWLEDGMENTS

This work was supported by MEXT KAKENHI Grant Number 22K04892.

## AUTHOR CONTRIBUTIONS

Conceptualization, A.N., Y.M., and T.M.; investigation – data collection, A.N. and H.K.; formal data analysis; A.N., Y.M., and T.M. data visualization, A.N.; writing – review & editing, A.N., Y.M., and T.M.; funding acquisition, T.M.; resources, T.M.; supervision, T.M.

## DATA AVAILABILITY

The data that support the findings of this study are available from the corresponding author upon reasonable request.

## SUPPLEMENTARY MATERIAL

## Supplementary Figures

**FIGURE S1.**
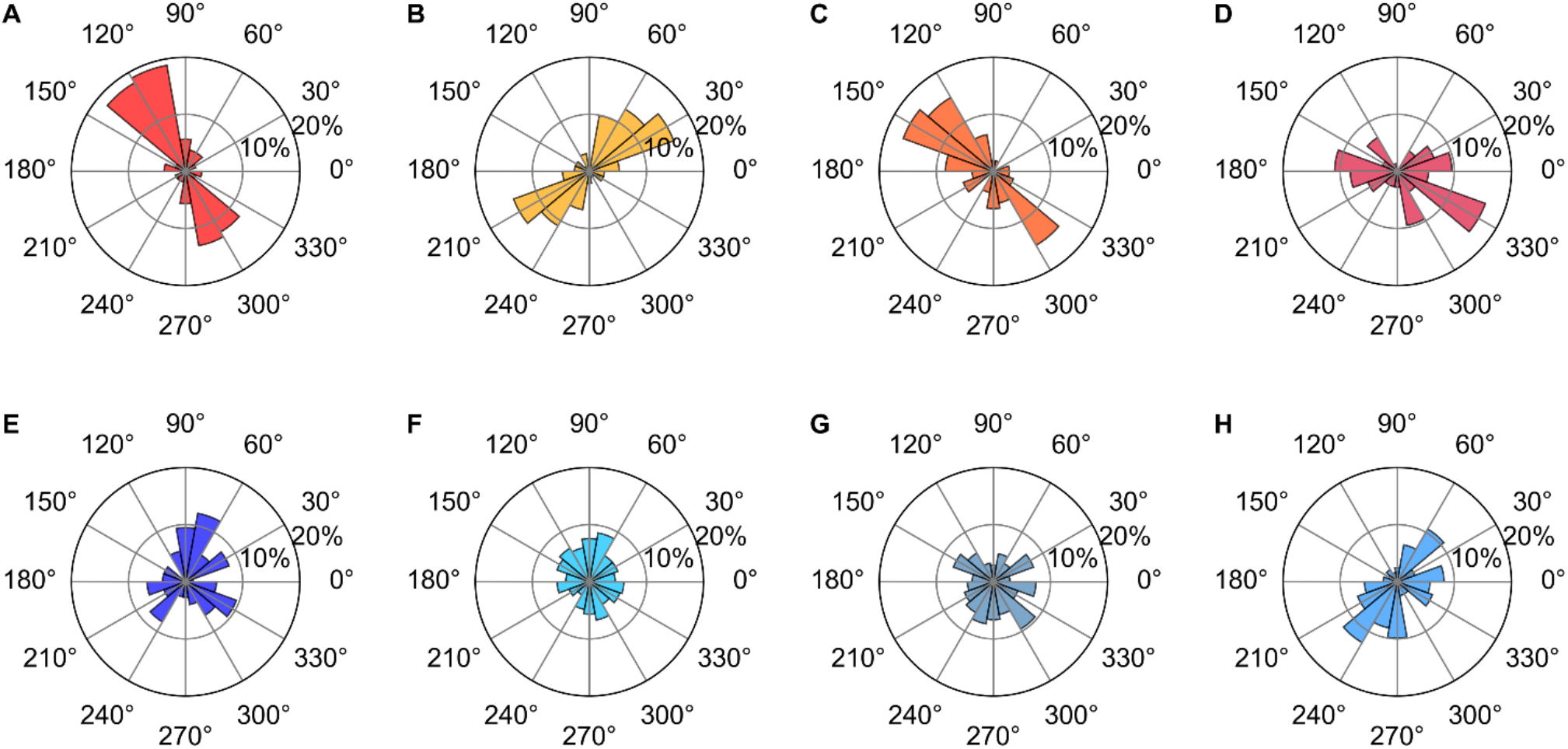
Angular distribution histograms of cell velocity for four isolated cells during elongation and retraction phases. (A–D) Histograms of cell velocity on 10 kPa PDMS: (A) n = 106, (B) n = 191, (C) n = 107, (D) n = 73. (E–H) Histograms of cell velocity on 400 kPa PDMS: (E) n = 74, (F) n = 265, (G) n = 241, (H) n = 122

**FIGURE S2.**
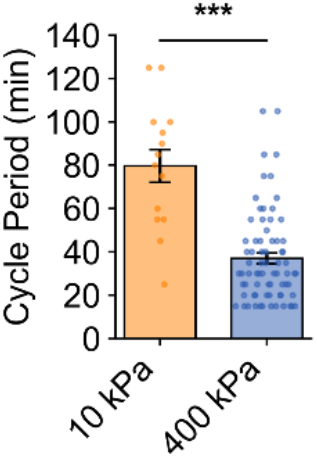
Motility cycle period of cardiac fibroblasts on 10 kPa and 400 kPa PDMS substrates. Cells on the 10 kPa substrate exhibited significantly longer motility cycles, 79.64 ± 7.53 minutes (n = 14), compared to 37.01 ± 2.47 minutes (n = 72) on the 400 kPa substrate. Data are presented as mean ± standard error.

## Supplementary Movies

**Movie S1**. Time-lapse movie of a cardiac fibroblast on 10 kPa PDMS substrate. The cell exhibits back- and-forth oscillatory movement over multiple motility cycles.

**Movie S2**. Time-lapse movie of a cardiac fibroblast on 400 kPa PDMS substrate. The cell exhibits random-like, non-directional movement over multiple motility cycles.

